# K-seq, an affordable, reliable, and open Klenow NGS-based genotyping technology

**DOI:** 10.1101/2020.11.16.360446

**Authors:** Peio Ziarsolo, Tomas Hasing, Rebeca Hilario, Victor Garcia-Carpintero, Jose Blanca, Aureliano Bombarely, Joaquin Cañizares

## Abstract

K-seq, a new genotyping methodology based on the amplification of genomic regions using two steps of Klenow amplification with short oligonucleotides, followed by standard PCR and Illumina sequencing, is presented. The protocol was accompanied by software developed to aid with primer set design. As the first examples, K-seq in species as diverse as tomato, dog and wheat was developed. K-seq provided genetic distances similar to those based on WGS in dogs. Experiments comparing K-seq and GBS in tomato showed similar genetic results, although K-seq had the advantage of finding more SNPs for the same number of Illumina reads. The technology reproducibility was tested with two independent runs of the tomato samples, and the correlation coefficient of the SNP coverages between samples was 0.8 and the genotype match was above 94%. K-seq also proved to be useful in polyploid species. The wheat samples generated specific markers for all subgenomes, and the SNPs generated from the diploid ancestors were located in the expected subgenome with accuracies greater than 80%. K-seq is an open, patent-unencumbered, easy-to-set-up, cost-effective and reliable technology ready to be used by any molecular biology laboratory without special equipment in many genetic studies.

## INTRODUCTION

The NGS revolution has induced a very rapid improvement in genotyping technologies; it is now standard to design experiments with hundreds of thousands of markers. This progress has accelerated gene and QTL analyses and allowed the popularisation of robust association studies as well as a marked increase in the scope of population diversity and evolution research in both model and non-model species. The most common genotyping approaches are shown in Table 1. Currently, no ideal universal technology is available for all experimental designs. The preferred platform depends on the number of samples, accuracy required, population structure, and cost and availability of each methodology for the target species.

**Table 1.** Common genotype methods

|  | Cost | RNA handling | Patented | Targeted | Predesign platform |
| --- | --- | --- | --- | --- | --- |
| WGS | High | No | No | Yes | No |
| RNAseq | High | Yes | No | No | No |
| Exome sequencing | High | No | No | No | Yes |
| Amplicon sequencing | High | No | No | Yes | Yes |
| GBS | Low | No | Yes | No | No |
| DArTseq | Low | No | Yes | No | No |
| rAmpSeq | Low | No | No | No | No |
| K-Seq | Low | No | No | No | No |
| SNP microarrays | Low | No | Yes | Yes | Yes |
| KASP | Low | No | Yes | Yes | Yes |
| SPET | Low | No | Yes | Yes | Yes |
| Seq-SNP | Low | No | Yes | Yes | Yes |
| Ion AmpliSeq technology | Low | No | Yes | Yes | Yes |

A popular approach is to target sets of genetic markers identified in previously resequenced populations. Several target SNP genotyping systems have been developed, such as microarray-based systems(1), KASP(2) (LGC group, UK), SPET(3, 4) (Nugen, CA, USA), and Seq-SNP in LGC. The genotyping cost per sample is usually low for these technologies, but they involve an initial costly SNP discovery and design step that must be paid upfront; so frequently, they are only available for model species. Moreover, few platforms are usually created for any given species due to the high prior development costs; thus, only some limited SNP sets tend to be available. A thoughtful initial SNP selection step is critical for the success of these platforms, but any preselected set of variants will have some drawbacks. Any particular experiment, for example, an F2 genetic map, will involve a restricted set of samples, so many of the preselected SNPs present in the genotyping platform will not be polymorphic and, thus, not useful at all. Moreover, although preselected sets of SNPs tend to work well for samples present in the populations that are sequenced during the design step, they can fail to capture the diversity of other populations. These sets of variants can also introduce diversity bias based on the populations involved during the platform design step, thus skewing the calculation of any population-level diversity and distance index.

Alternatively, all samples could be genotyped by resequencing. This approach has the advantage of not requiring any prior information, so it can produce unbiased sets of variants, and there is no initial cost associated with developing a marker platform. Many genome-wide association studies and population history analyses have used Whole Genome Sequencing (WGS). Unfortunately, WGS at a high coverage is still expensive, particularly for species with large genomes; additionally, for many studies, WGS is unnecessary. Costs can be reduced by sequencing at a low coverage, but this cheaper approach usually involves an imputation step that complicates posterior bioinformatic analysis, and its success depends on the genetic structure of the populations analysed. Consequently, several Genomic Reduced Representation (GRR) technologies have been developed. These methodologies are based on the reproducible sampling of genomic regions across multiple samples before the sequencing step and usually generate unbiased sets of variants from high-coverage sequencing at a low cost.

GRR technologies differ in which and how genomic regions are targeted. A simple way to select transcribed regions is to use RNA-seq. The transcriptome requires fewer reads than the genome to be characterised, so it is cheaper than WGS. One RNA-seq drawback is that it forces the researcher to handle RNA instead of the more resilient DNA, possibly creating problems, for example, in field studies. Moreover, RNAseq libraries are usually more expensive to generate than those required in other approaches, such as GBS(5) and RAD-seq(6, 7). Exome capture(8) and other amplicon-based genomic selection technologies, such as Ion AmpliSeq technology (Thermo Fisher Scientific, Waltham, MA), have been developed as alternatives to RNA-seq. These are effective ways to select particular genomic regions, but they require a complex design and usually involve higher costs.

Multiple restriction enzyme-based genomic selection technologies, such as GBS, RAD-seq or DArTseq™(9, 10), have been developed. The cost of these methodologies is usually much lower than exome capture or RNA-seq. The cost by sample is similar to that of the marker set platforms, without considering the platform development cost. These restriction-based approaches mainly differ in the enzymes and purification protocols used. GBS is one of the simplest and most popular choice. In this case, the fragments are selected by digesting genomic DNA with one or two enzymes, usually *ApeK*I or *Pst*I. Next, adapters are ligated to the fragments of DNA prior to PCR amplification. GBS does not involve any purification or size-selection step, and the amplified fragments are directly sequenced using Illumina. The popularity of GBS is due to its ability to select a small fraction of the genome, thus allowing the researcher to genotype at both a high coverage and a low cost. The number of markers obtained will depend on the restriction enzyme(s) used, genome size, and variability across samples.

These restriction enzyme-based technologies only require a pilot assay to select the appropriate enzyme, so they are cheap to establish in any new species and produce very affordable genotypes. They represent the common approach in non-model species and small-to medium-scale projects (those with hundreds of samples). Moreover, they are not limited to a previously selected set of SNPs, so they are not subject to target marker set platform biases. A drawback of these non-targeted genomic selection protocols is that the researcher cannot choose specific regions, such as genes of interest, to be genotyped. Another minor shortcoming is that they require more bioinformatic expertise than the target marker platforms, although this support is usually provided by the sequencing provider. However, another practical limitation is that many of these technologies are patented, requiring a license to be used. This requirement increases the cost and limits the number of service providers available.

GRR technologies can be improved, and new platforms are still being developed(11–15). It would be ideal to have open, patent-unencumbered technologies that could be easily used by any molecular biology laboratory. An example of a GRR methodology not based on enzyme restriction is rAmpSeq(13), which uses primers located in repetitive sequences to amplify middle repetitive regions. In this study, we present such a technology, K-seq. The protocol is based on the selection of thousands of genomic fragments using two amplification cycles with Klenow polymerase and short primers, followed by PCR amplification. Through systematic selection of short primers, K-seq targets fragments located on nonrepetitive regions that are easily mappable on downstream analyses. The combination of Klenow polymerase and short oligonucleotides provided both good reproducibility and thousands of genotyped SNPs at a very low cost in the tests carried out in Solanaceae, wheat and dogs.

## MATERIAL AND METHODS

### Primer selection

The short oligonucleotides used for the different species were selected by using a software application developed for this purpose, Primer Explorer (https://github.com/bioinfcomav/primer_explorer2). The algorithm used for the oligonucleotide selection process is shown in supplemental data file 1. Briefly, only k-mers with GC contents between 35% and 75% were considered. The 1,000 most abundant k-mers in the regions considered to be nonrepetitive in the genome annotation were selected. Those were combined to create potential oligonucleotide sets comprising 10 k-mers that were further analysed to check for compatibility in a PCR. In these sets, the oligonucleotides were split into forward and reverse primers because, as described below, the K-seq protocol amplifies and sequences only forward-reverse combinations but not forward-forward or reverse-reverse combinations. Virtual PCR was carried out for the oligonucleotide sets deemed compatible according to Primer3 2.4.0(16). Based on the results of this PCR, a report that included the number of predicted single and repetitive products for each primer pair within each oligonucleotide set was generated. The final oligonucleotide set chosen for each species was based on the report that included the predicted number of products for each primer pair within the 10 primer oligonucleotide set.

### Biological materials and DNA isolation

The sample passport data are shown in Supplemental Table 1. Solanaceae and wheat DNAs were isolated from young leaves using a CTAB-based protocol(17). Dog DNA was isolated from clinical blood samples provided by CEU Cardenal Herrera Veterinary Hospital of University of Valencia using the Wizard Genome DNA Purification Kit (Promega, WI, USA). The use of dog blood samples was approved by the Animal Experimentation Ethics Committee of CEU Cardenal Herrera University of Valencia. After extraction, the DNA was checked by agarose gel electrophoresis and quantified with Qubit (Thermo Fisher Scientific Inc. MA, USA).

### Database sequences

Additional publicly available dog genome sequences were downloaded from NCBI. The breed and genomic information for each sample can be found in Supplemental Table 2. The tomato GBS sequences are available in NCBI, Bioproject PRJNA644494 (Supplemental Table 3).

### K-seq genotyping

For the Solanaceae and dog reactions, 200 ng of DNA was used; for wheat, 600 ng was employed. The detailed K-seq protocol is described in Supplemental Data File 2 and Figure 1. Briefly, genomic DNA was denatured and annealed at 37°C using forward primers. The primers used for every species are described in supplemental data file 3. After forward primer annealing, the DNA Pol I large Klenow fragment (New England Biolabs, MA. USA) was used in the first DNA synthesis step. Once the polymerase was inactivated at 75°C, the remaining single-chain DNA, which included uncopied genomic DNA and free forward primers, was destroyed by incubating the reactions with Exonuclease I at 37°C (New England Biolabs, MA. USA). After exonuclease inactivation at 80°C, both a second Klenow denaturation and amplification cycle followed, this time using reverse primers. Finally, the first two amplification cycles were ended by a further Exonuclease I treatment. This exonuclease-treated Klenow amplification was then used as a template in a standard 15-cycle PCR (Taq 2X Master Mix; New England Biolabs, MA. USA) using Illumina IDT-NXT primers carrying dual indexes (Supplemental Data 4).

**Figure 1.**
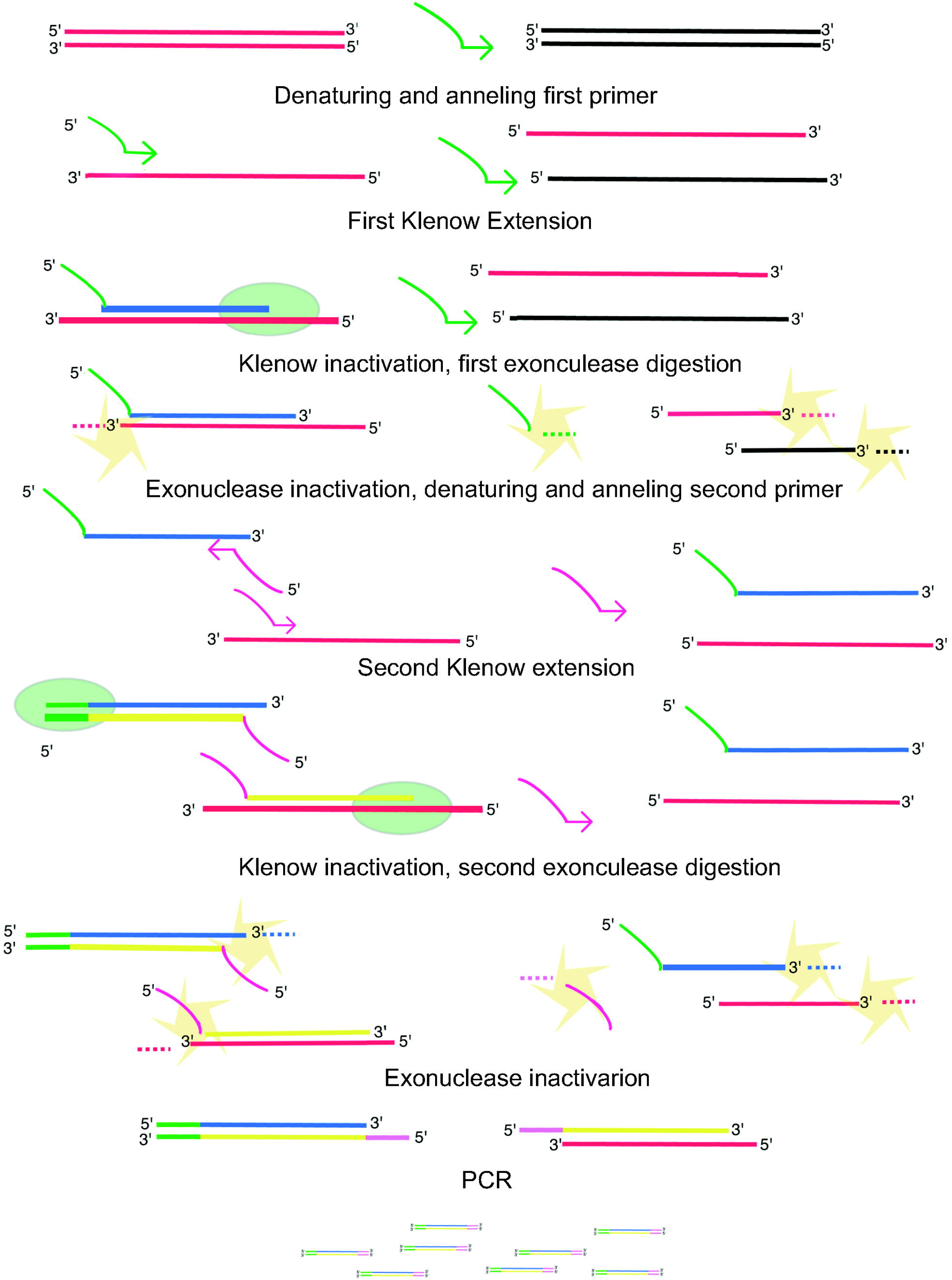
K-seq protocol

The number of cycles for the PCR, 15, was chosen by carrying tests with different cycle numbers. The objective was to perform sufficient cycles to amplify the sample to be sequenced using Illumina while avoiding overamplification that could introduce artefacts. The PCR products were run in agarose gels, and we selected 15 cycles because they did not create artefactual long DNA products.

To check the reproducibility, the tomato control samples were processed in two completely independent experiments. For sequencing, all the samples were pooled in three libraries. The first comprised dog samples, the second comprised wheat samples and, the third included Solanaceae samples. All the pools were built by mixing equal volumes of each PCR product. In future experiments, to equalise the final number of reads obtained for each sample, it might be better to quantify the DNA present in each PCR before pooling them. The three libraries were size-selected and sequenced together in a HiSeq 2500 lane (Illumina, CA, USA) by CNAG-CRG (Barcelona, Spain). The reads obtained are available in NCBI bioproject PRJNA635525.

### Mapping and SNP calling

The possible adapter sequences present in the Illumina reads were removed using Cutadapt 1.18(13). The trimmed reads were mapped against the reference genome (Supplemental Table 1) using BWA-MEM 0.7.17-r1188(18), and only the mapped reads with the highest MAPQ, 56, were used for Freebayes v1.3.1-19-g54bf409(19) SNP calling. The Freebayes parameters used were min_mapq=57, min_base_quality=20, min_coverage=10, and min_alternate_count=0.1. For the analyses, only the highest quality SNPs were used: the genotypes with a coverage lower than 10 were set to missing, and the SNPs with a missing genotype rate above 0.6 were removed.

### Genetic analysis

The mapping statistics were extracted using Samtools 1.9 (20) stats. SNP stats were performed using homemade scripts (https://github.com/bioinfcomav/variation5/) for Solanaceae and dog and wheat with rtg 3.10.1 (https://www.realtimegenomics.com/products/rtg-tools) for wheat. Genetic distances and Principal Coordinate Analyses (PCoA) were calculated using the Python library Variation5 (https://github.com/bioinfcomav/variation5/). PCoA was carried out using SNPs with a missing genotype rate under 0.05.

## RESULTS

### Oligonucleotide selection and K-seq library construction

Oligonucleotide combinations used for Solanaceae, dog and wheat were chosen considering the reports created by Primer Explorer software for the tomato, dog and wheat reference genomes (supplementary data file 4). The oligonucleotide size used was 9 bp for the species with larger genomes—dogs and wheat—and 8 bp for tomato. For all species, the final primer combination comprised three oligonucleotides (supplemental data file 4). Primer Explorer predicted 6,541, 12,134, and 12,119 products, that could be sequenced, for tomato, wheat and dog, respectively. The selection process chose oligonucleotides from k-mers enriched in single-copy genomic regions, so the primer combinations should generate less repetitive products than those created from random k-mers. The putative numbers of mapping fragments in nonrepetitive regions for these species were 4,187 in tomato, 11,286 in wheat and 7,150 in dogs.

Genomic DNA fragments were amplified following the protocol explained in the Materials and Methods: two rounds of Klenow polymerase amplification followed by PCR. The tomato results are shown in Supplemental Figure 1 A as an example of the fragment size distribution obtained. Finally, the libraries were combined into species-specific pools that were size-selected (300 to 700 bp) before sequencing in a HiSeq2500 Illumina lane (Supplemental Figure 1 B).

### Mapping and variant calling

The number of reads obtained per sample ranged from 0.8 to 50 million (Table 2, Supplemental Table 5). This high variability might be due to the lack of DNA quantification of the PCR-amplified products before library pooling; thus, the reaction products should be quantified before pooling. The processed reads were mapped against the reference genomes (Supplemental Table 5) with average mapping rates of 92.7%, 94.0% and 95.9% for tomato, wheat and dog, respectively. After filtering out reads with MAPQ lower than 56, the percentages of the remaining reads were 71.2%, 36.2% and 89.9%. The percentage of univocally mapped reads was much lower in wheat, representing an expected limitation due to wheat’s allohexaploid nature. The size distribution of the mapped fragments (Supplemental Figure 2) was very similar to that of the sequenced libraries (Supplemental Figure 1 B). The density of the mapped fragments along the genome for tomato, wheat and dog was uniform with very few regions with higher or lower density in some chromosomes (Supplemental Figure 3).

**Table 2.**
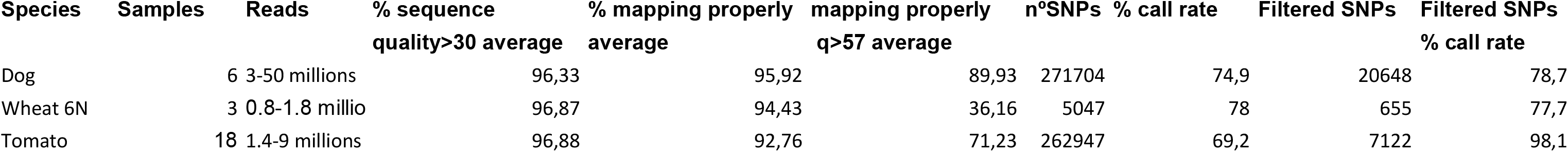
Read, mapping and SNP statistics

Variants were called by Freebayes, and sample statistics are shown in Supplemental Figure 4. The mean genotype call rates for the high-quality variants (Table 2) were 98.8%, 77.7%, and 78.7% for tomato, wheat and dogs, respectively. The wheat and dog reactions generated a highly variable number of reads (Supplemental Figure 4), that, in turn, accounted for the decreased wheat and dog mean genotype call rates. The observed heterozygosity varied from 13% to 35% in the dog samples and from 8% to 18% in the wheat samples. However, in tomato, as expected, it was much higher (58%) in the controls built by mixing isomolar quantities of DNA extracted from two samples, Heinz1706 and LA1589, originating from two different species: *Solanum lycopersicum* and *Solanum pimpinellifolium*.

### Reproducibility

Tomato samples were genotyped twice in two independent reactions. To compare both experiments, 1.7 million reads were selected randomly from each sample (1.7 million was the number of reads obtained for the sample with fewer sequences), and the coverages obtained in the two independent replicates were compared. The coverage correlation coefficients were 0.84 for Heinz1706, 0.73 for LA1589, and 0.86 for the isomolar mix Heinz1706+LA1589. The percentage of SNPs with matching genotypes in the two replicates was 98.54% for Heinz1706, 98.35% for LA1589 and 93.58% for the mixed sample Heinz1706+LA1589. In all cases, the non-matching genotypes were due to positions deemed heterozygote in one replicate but homozygote in the other. Of the 6,624 SNPs found in these samples, 5,682 (86%) were genotyped in the three samples. Additionally, given that the sample Heinz1706+LA1589 was created by an equimolar mix of the other two, it was possible to calculate how many of the SNPs expected to be heterozygous in this mixed were truly heterozygotes according to the SNP caller: 98%.

### Comparative analysis of K-seq and GBS

GBS genotypes were available for all three tomato samples. For each of these samples, Heinz1706, LA1589 and Heinz1706+LA1589, two independent GBS and two K-seq replicates were analysed by selecting 1.7 million reads from each sequencing reaction. The mean percentage of mapped reads was similar for GBS and K-seq, 92% and 95%, respectively. However, after filtering these mapped reads according to their MAPQ value, to remove repetitive regions, the number of mapped reads was higher for K-seq than for GBS, with averages of 39% and 70%, respectively (Figure 2A). The final number of filtered SNPs was 7,701 for K-seq and 3,062 for GBS. Moreover, for any minimum coverage that one would care to select, the number of SNPs obtained was always higher for K-seq than for GBS (Figure 2B). The correlation of SNP coverages between replicates was 0.81 for K-seq and 0.70 for GBS. The percentage of matching genotypes between replicates ranged from 93% to 99% for different samples and was similar for K-seq and GBS (Figure 2C). Mismatched genotypes were more common in the Heinz1706+LA1589 samples.

**Figure 2.**
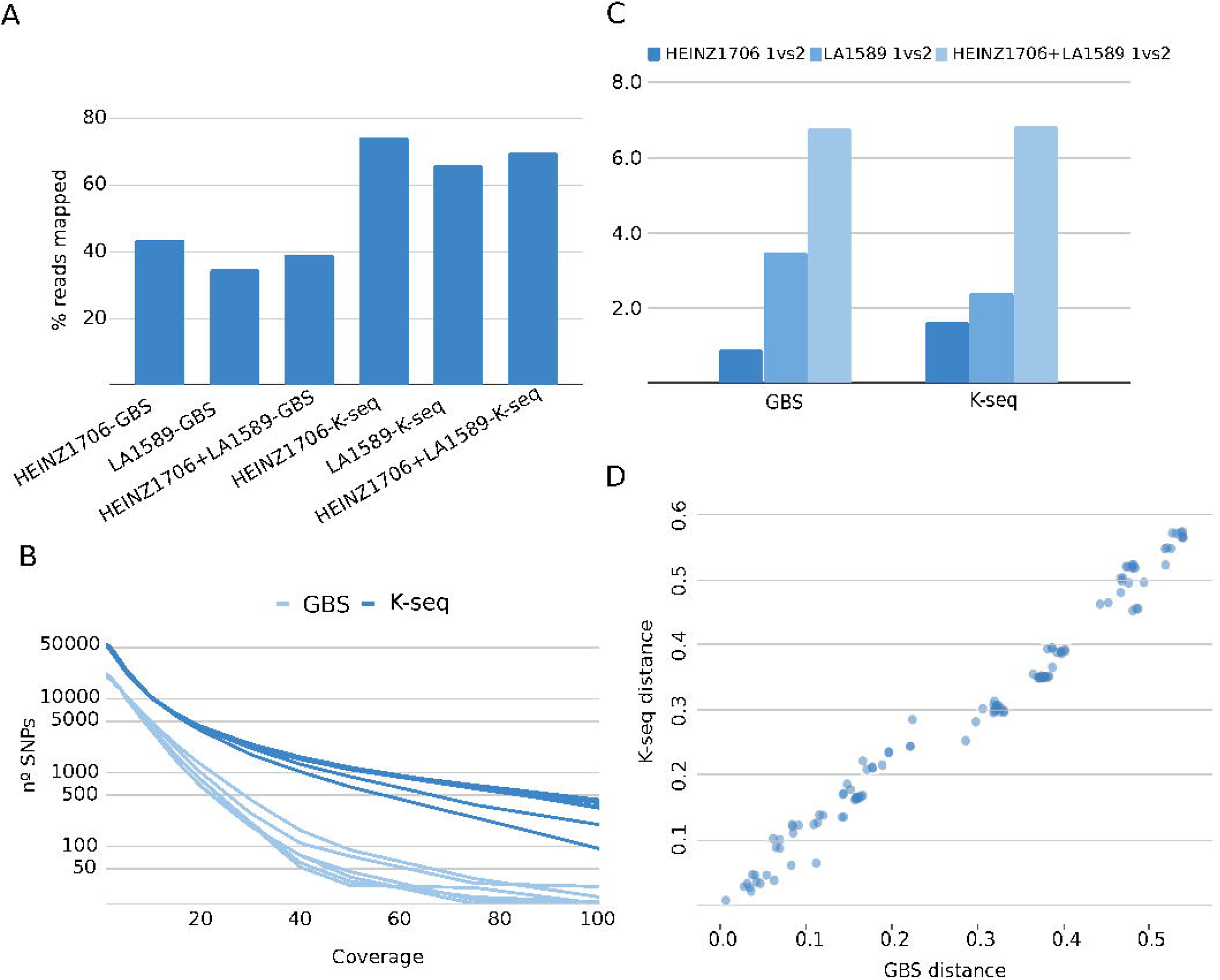
GBS and K-seq comparative. A, Percentage of mapping reads. B, Coverage distribution of genotyped SNPs. C, Percentage of genotype differences between replicates. D, Correlation between genetic distances calculated with GBS and K-seq.

Additionally, we analysed 19 tomato samples that were genotyped by K-seq and GBS. The correlation between the genetic distances calculated among samples from K-seq and GBS genotypes was 0.99 (Figure 2D).

### Comparison of the predicted and sequenced regions

The genomic locations that were predicted using Primer Explorer were compared with the amplified fragments. The difference between the prediction and experiment was substantial. Of the regions sequenced using a coverage higher than 10X, only 2% to 4% had been previously predicted by the software (Supplemental Table 4). The main difference between prediction and experiment was due to the sequencing of many unexpected fragments. This effect was mainly due to Klenow polymerase amplifying regions using primers with one or more mismatches to the corresponding genomic template. The percentage of mismatched primers varied from 23% to 44% depending on the species and primer. In future experiments, longer oligonucleotides or modified Klenow annealing conditions could be tested to reduce this effect. Alternatively, to improve the prediction accuracy, these mismatches could be included in future versions of Primer Explorer. Despite these shortcomings, the number of genes covered by the prediction and number of genes finally sequenced were similar (Figure 3); moreover, the oligonucleotide set and Klenow annealing conditions are likely to remain stable in any species once an assay has been successful, so Primer Explorer will only be used once for any given species in most cases.

**Figure 3.**
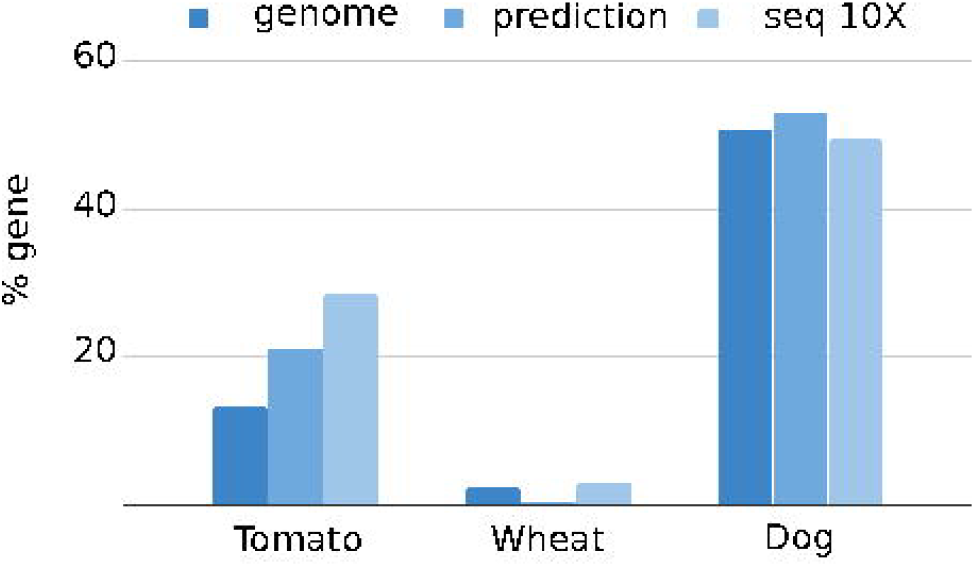
Percentage of bases located on genes in the genome, predicted fragments and bases sequenced with a coverage larger than 10X

### Performing K-seq in related species reusing oligonucleotide sets

To test whether genotyping a species without a reference genome is possible using K-seq, we used the tomato-derived oligonucleotide set in other Solanaceae. The sequences obtained were mapped to the respective genomes (Figure 4). The process generated good genotypes in all species tested: potato, pepper, eggplant, and even in the more distant Petunia × hybrida. The numbers of SNPs obtained using the tomato oligonucleotide set were 29,090 in potato, 3,316 in pepper, 2,263 in eggplant and 4,389 in Petunia x hybrida.

**Figure 4.**
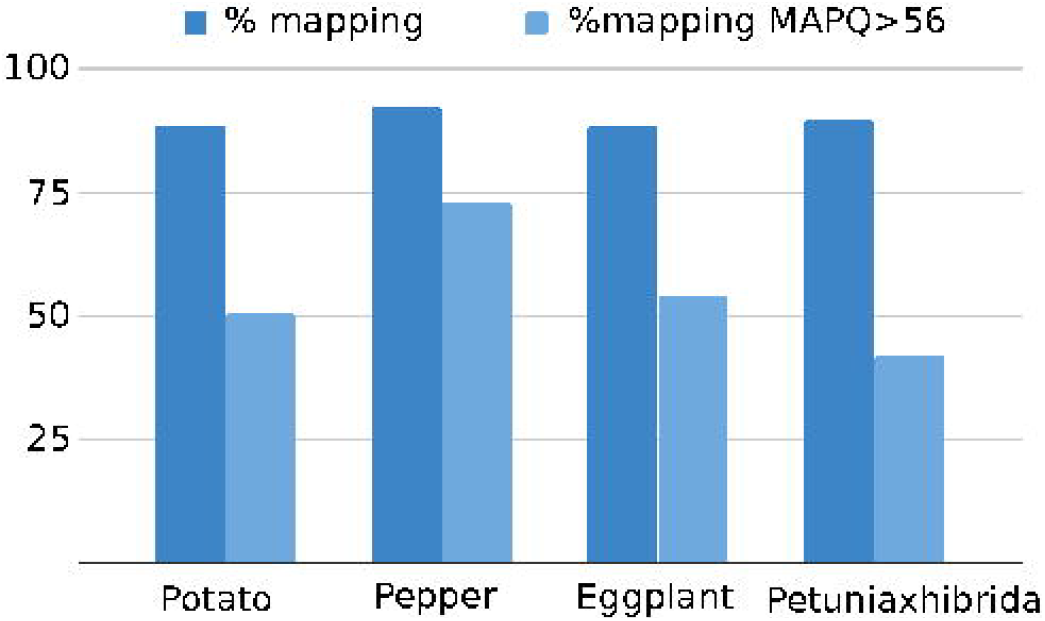
Mapping statistics of the Solanaceae samples genotyped with tomato short primers

### K-seq in mammals

We also wanted to test the K-seq technique in species with very different genome architectures; thus, we amplified and sequenced six dog samples as mammalian representatives. The mapping statistics showed that K-seq performed as expected (Table 2). Moreover, the oligonucleotides were selected by Primer Explorer to be located preferentially in single-copy regions; once the reads were mapped, 49.23% of the 10X or higher coverage regions were found within genes. Additionally, to test the performance of K-seq as a possible tool for animal population genetic studies, we compared some of the dog WGSs publicly available using our K-seq genotyping. PCoA including both the WGS and K-seq samples was carried out (Figure 5). In general, no clear bias was observed between the K-seq and WGS in the PCoA; moreover, the Yorkshire terrier and Poodle breeds were genotyped by both technologies, and their respective samples were found at short distances in the PCoA.

**Figure 5.**
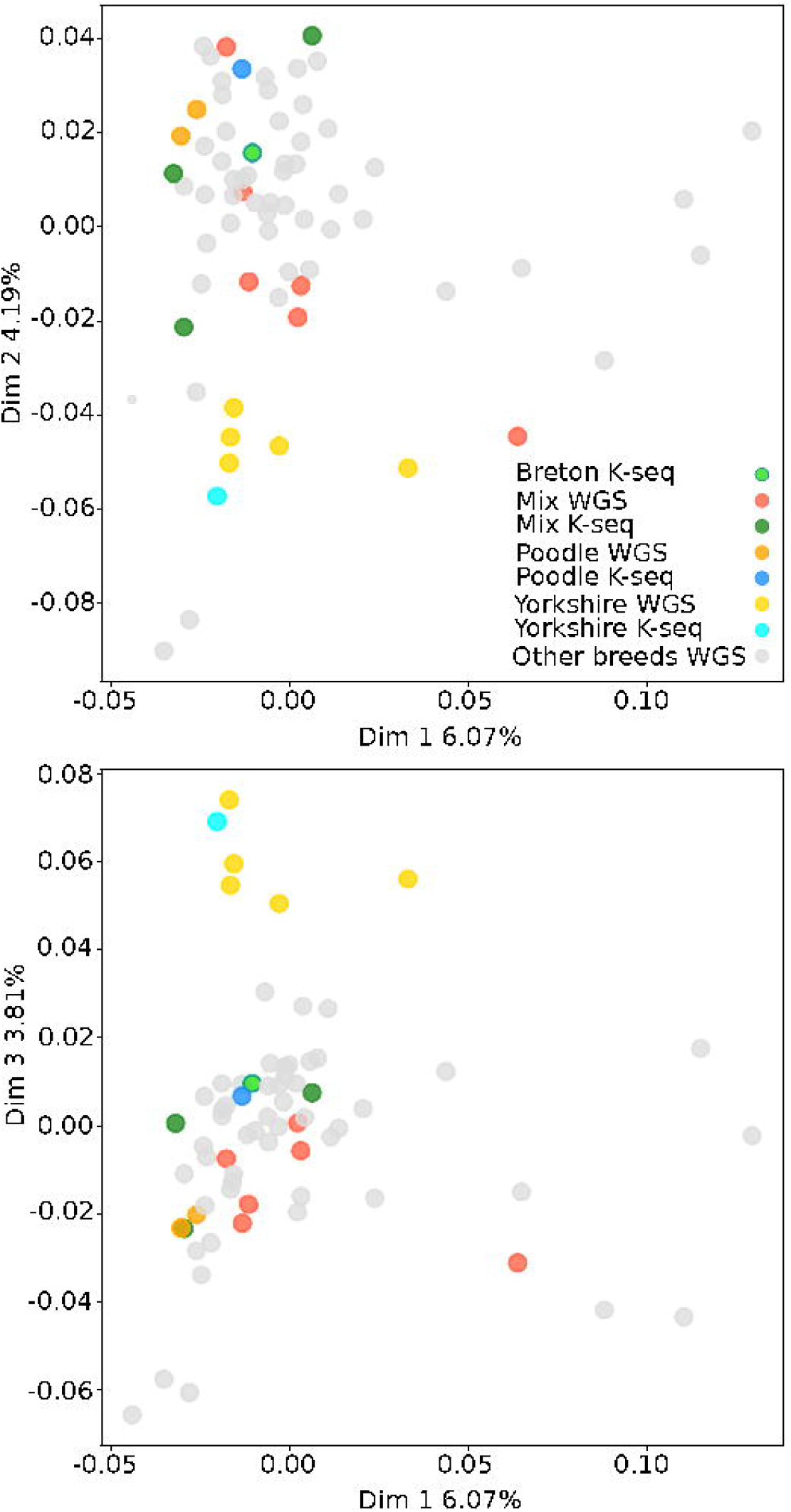
PCoAs of WGS and K-seq dog samples.

### Genotyping individuals with different ploidies

Finally, we also wanted to check the K-seq performance in a species complex with different ploidies. Primer Explorer was used to analyse the Chinese Spring v1.0 hexaploid maize reference genome, and the oligonucleotide set selected was used to genotype the species *Triticum aestivum* subsp. *vulgare* (6x)*, Triticum aestivum* subsp. *speltoides* (6x)*, Triticum turgidum* subsp. *dicoccon* (4x)*, Triticum turgidum* subsp. *turgidum* (4x)*, Triticum turgidum* subsp. *durum* (4x)*, Triticum urartu* (2x)*, Triticum monococum* subsp. *boeoticum* (2x)*, Aegilops speltoides* (2x) and *Aegilops tauschii* (2x). The percentage of mapped reads to the reference wheat genome varied with the sample ploidy (Figure 6A). The analysis showed that the reads mapped with high MAPQ were subgenomic specific (Figure 6B). The reads originated from the diploid, tetraploid and hexaploid samples mapped to the expected subgenome, and K-seq could identify specific SNPs for every chromosome. The number of genotyped SNPs varied with the read number, ploidy and variability of each sample. In the hexaploid samples (1 to 2 million reads), 4,000-5,000 SNPs were genotyped at a 10X coverage; in the tetraploid sample Svevo (9 million reads), 22,009 SNPs were obtained; and in the diploid sample BGE18908 (11 million reads), 6,147 SNPs were generated.

**Figure 6.**
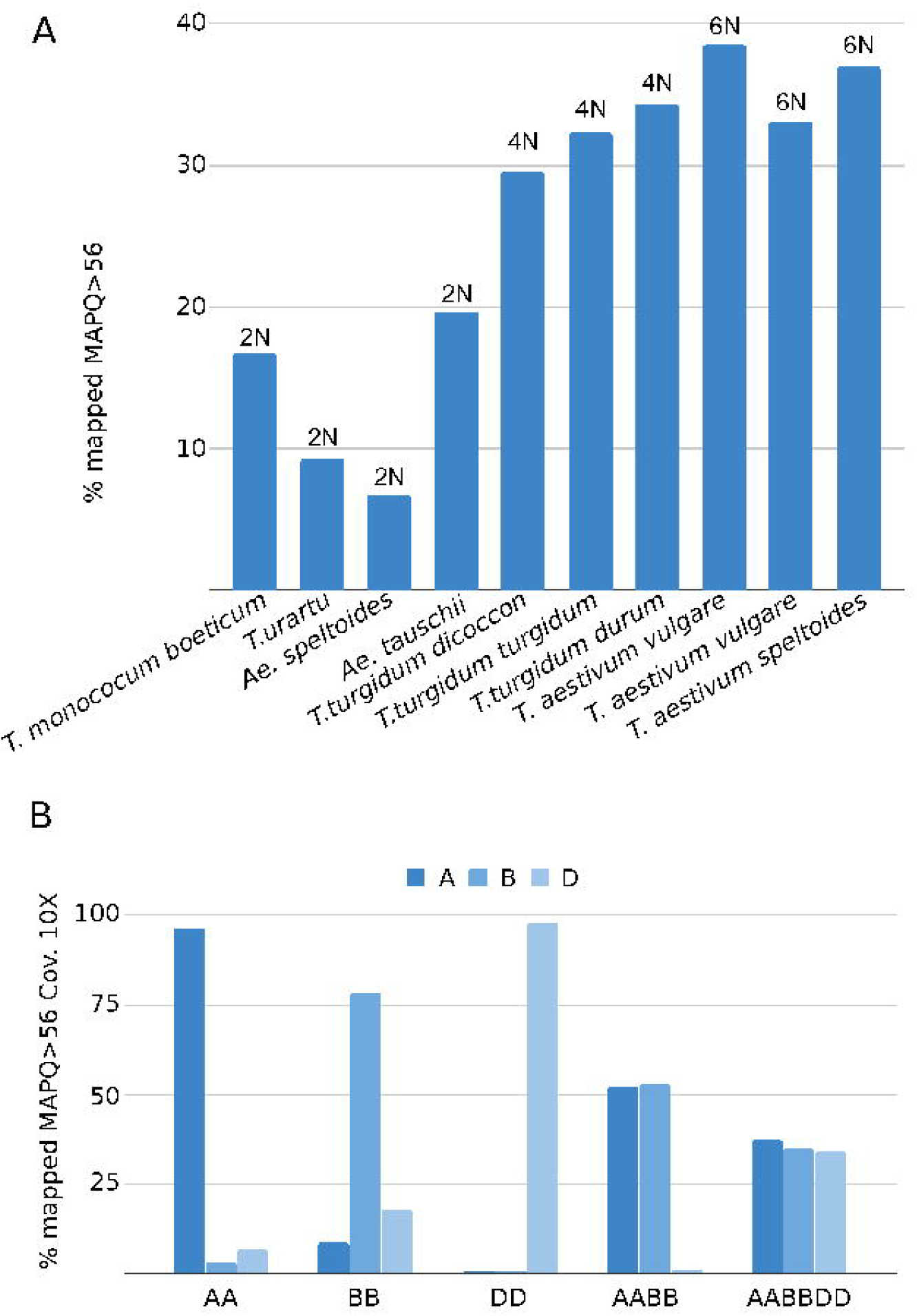
A, Percentage of mapping reads with MAPQ larger than 56 in the genome of allohexaploid wheat. B, Percentage average of positions with a coverage greater than 10X using mapping reads with MAPQ greater than 56 in the different subgenomes of the diploid and polyploid samples.

## DISCUSSION

K-seq is a reliable and affordable genotyping technology, and Klenow-amplified fragments are suitable for Illumina sequencing. K-seq has been successfully used to genotype species as diverse as tomato, petunia, wheat and dogs. We identified 262,947, 5,047 and 271,704 SNPs in the tomato, wheat and dog experiments, respectively. K-seq also managed to genotype 20,090 SNPs in a tetraploid potato using the tomato primer set. Moreover, in tomato, K-seq was compared with GBS and provided very similar genetic distances between samples; in dogs, no bias was detected between K-seq and WGS samples. Additionally, in our laboratory, we used K-seq to genotype an F2 population of 151 plants, and we built a genetic map with 147,326 SNP markers with 2.4 million reads per F2 individual (manuscript in preparation).

K-seq is very versatile, and the number of fragments amplified and sequenced can be easily modified by adding or removing compatible primers from the primer set, which can be combined with the number of Illumina reads ordered. For instance, we could use longer oligonucleotides or primer sets with fewer primers to reduce the number of sequenced fragments. Thus, we would cover fewer genomic regions that would require fewer reads for the same coverage.

K-seq also genotyped species with large polyploid genomes; thus, an experiment with diploid, allotetraploid and allohexaploid wheat was carried out. As expected, using the tetraploid and hexaploid species, fewer reads (approximately 30%) were mapped against the hexaploid wheat genome reference with very high (56) MAQP values. However, from those, highly specific markers were developed and tested by mapping the diploid species reads that corresponded to the different allohexaploid subgenomes: A, B and D. For *Triticum urartu* (AA) and *Triticum monococum* (AA), 98% and 94% of the reads corresponded to subgenome A, respectively. For *Aegilops speltoides* (BB), 78% corresponded to B; for *Aegilops tauschii* (DD), 98% corresponded to D. Thus, it is possible to obtain specific genetic markers using K-seq, even in species with a genome as complex as wheat.

With 1 to 2 million reads per sample and setting a highly strict MAPQ filter, 5,047 K-seq SNPs were identified on the three hexaploid samples. By lowering the MAPQ threshold to 39, 9,754 SNPs were obtained from the same reads. This result is comparable with previous DArTseq and GBS wheat studies. For example, Riaz and coworkers(21) identified 34,311 DArTseq markers in 295 samples, and Chu et al.(22) obtained a similar result using GBS in 378 samples. Finally, Vikram and collaborators(23) obtained 20,526 SNPs in 8,416 samples.

K-seq was reliable, and the genomic sites sequenced were highly correlated between the two tomato experiments. When genomic sites with coverages higher than 5X were considered, the correlation between the coverages obtained in the two replicates was 0.8. Genomic regions sequenced in different samples were also very similar. The mean call rate for the different experiments ranged from 69.2% to 78%; thus, most of the unfiltered SNPs were genotyped for all samples in every experiment. K-seq also detected heterozygous genotypes in the mixed control tomato sample with 98% accuracy.

In the dog and tomato experiments, the K-seq, WGS and GBS results were successfully compared. The results obtained for the different technologies were very similar; thus, no biases between them were detected in these assays.

K-seq uses short oligonucleotides, similar to RAPDs(24), another technology that also uses short oligonucleotides but is notorious for its lack of reproducibility. However, RAPDs are based on standard Taq PCR amplification, and the annealing steps are carried out at temperatures above 50°C in which the short primers do not hybridise reliably. The key difference between RAPD and K-seq is the use of Klenow polymerase, an enzyme that operates at 37°C, a temperature more suitable for the hybridisation of short oligonucleotides, improving the reproducibility. The drawback, of course, is that Klenow is not thermostable, but the primers include a long adapter sequence that is also compatible with Illumina sequencing; thus, after the first two amplification cycles, standard Taq-based PCR can be used.

For genotyping technologies cost is a critical issue to consider. The cost per sample of a K-seq reaction, excluding sequencing, is approximately 8€. Klenow is the most expensive reactive, and laboratories buying it in bulk could probably manage to reduce the price. Moreover, the reaction could be adapted to low reaction volumes and automated with standard liquid handling robots to further reduce the cost. This protocol could also be adapted to other low optimal temperature DNA polymerases. Another advantage of the protocol presented is that size selection is performed once the libraries are pooled; thus, the protocol can be performed by the sequencing service and for only one pooled library. Therefore, we have developed a genotyping technology easy to implement in a small laboratory with basic molecular genetic infrastructure that does not require any previous genomic knowledge and that is comparable in cost to GBS.

To date, GBS has been a great option to genotype a high number of SNPs at a low cost. GBS has been a great success in non-model species. It has been extensively used in genetic analysis, population genetics, and breeding in a wide range of species. The results obtained with K-seq were better than those obtained with GBS. In our tomato experiments, K-seq was even more efficient at obtaining high-quality SNPs than GBS because the K-seq-amplified fragments tended to be located in single-copy regions with a higher frequency than the GBS fragments: 70% vs 40%. Regarding GBS, the restriction enzyme of choice is ApeKI, precisely because its cut sites are enriched in the gene-dense genomic regions; however, even when that precaution is considered, K-seq performs better. Moreover, the cost of GBS genotyping has recently increased, and the number of sequencing services offering the technology has been reduced because of the licencing policies of GBS patent holders. K-seq is open and offered free of patenting encumbrances; thus, it can be freely used.

Moreover, K-seq is highly versatile, and the genomic regions amplified can be changed by adding or removing oligonucleotides from the primer set using Primer Explorer reports. Although Primer Explorer cannot predict with precision the exact genomic fragments amplified experimentally, its prediction has been sufficient to locate k-mers enriched in gene-rich single-copy genomic regions. Moreover, given that the analyses of the sequences obtained allowed us to determine that the main problem with the prediction was small mismatches between the primers and genomic DNA template, in the future, the Klenow annealing conditions could be made more stringent or the software could be modified to account for these mismatches.

K-seq could be used even in species with no previous genome draft available. One option would be to perform prelaminar Illumina sequencing to determine the species k-mer composition, but that process has been proven to be not required. The Solanaceae data showed that the primers designed using the tomato genome reference could be successfully used to genotype a species as distant as petunia. Thus, K-seq can be used in species with no previous genomic information using primers designed for another species of the same genus. To perform a more accurate analysis, in the current study, we used the available genome sequences of all the Solanaceae studied. Of course, this would not be possible in a species with no genome assembly. For those cases, freely available open-source bioinformatic tools have been developed, such as Tassel(25) or Stacks (https://catchenlab.life.illinois.edu/stacks/), which could be adapted to analyse K-seq data.

In summary, we have described K-seq, a low-cost new genotyping technology that is easy to use by most laboratories, adaptable to any species and not encumbered by patents. We hope that K-seq will be useful in future genetic studies and can be improved even further by the community to obtain better results at lower costs.

## AVAILABILITY

The Primer-explorer2 and Variation5 applications are available in the GitHub repository https://github.com/bioinfcomav

## ACCESSION NUMBERS

The reads are included in NCBI’s bioproject PRJNA635525.

## SUPPLEMENTARY DATA

Supplementary Data are available at NAR online.

## ACKNOWLEDGMENTS

We thank Dr. Francisco Vázquez and Dr. Laura Pascual of University Polytechnic of Madrid for providing wheat seeds and the COMAV GenBank for providing tomato seeds. We also thank Dr. Begoña Ballester and the clinical analysis laboratory of the Clinical Veterinary Hospital of CEU Cardenal Herrera University for providing dog blood samples. We are grateful to the CNAG-CRAG and Dr. Úrsula Estada of SCSIE-UCIM of the University of Valencia for their help in the purification libraries and sequencing. Finally, we want to thank Esther Cañizares her help with images design.

