## supplemental files for "K-seq, an affordable, reliable, and open Klenow NGS-based genotyping technology"

Bionalyzer tomato sample

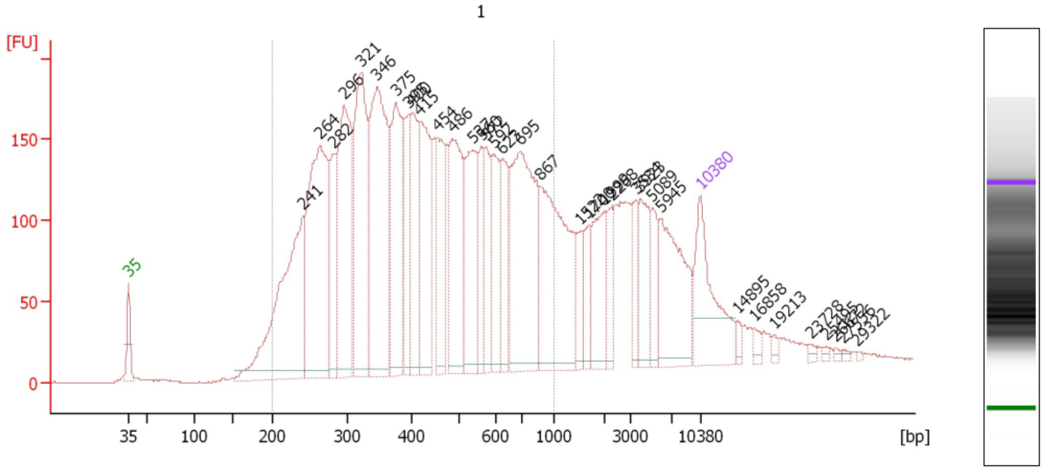

Bionalyzer size selected tomato sample

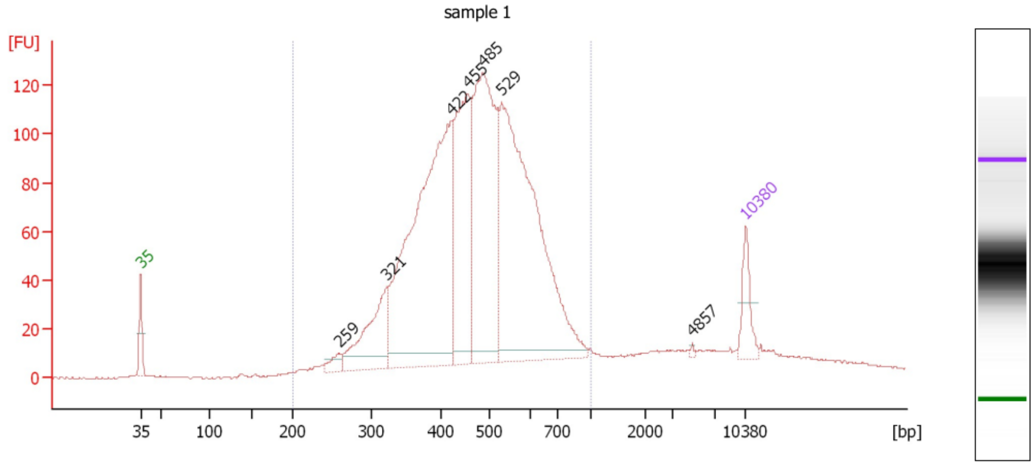

SFigure1. Bionalyzer results of tomato pool  sample

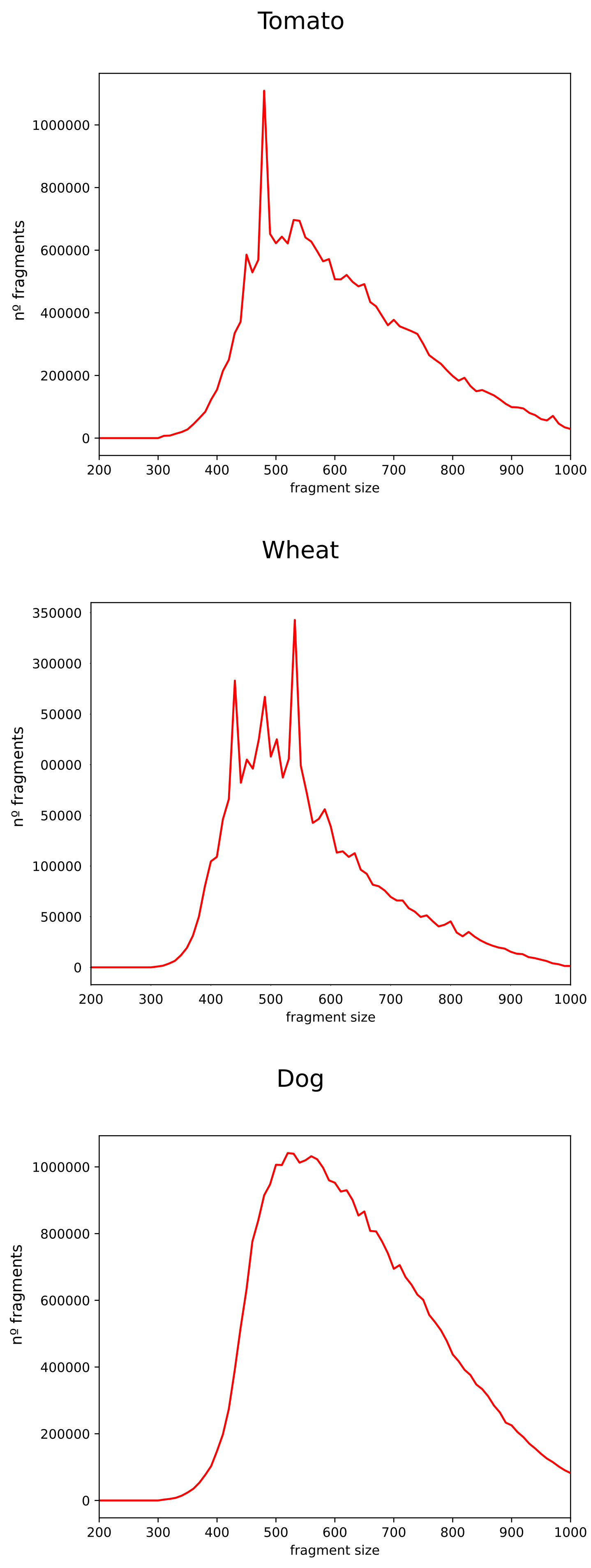

SFigure 2. Size distribution of mapped fragments

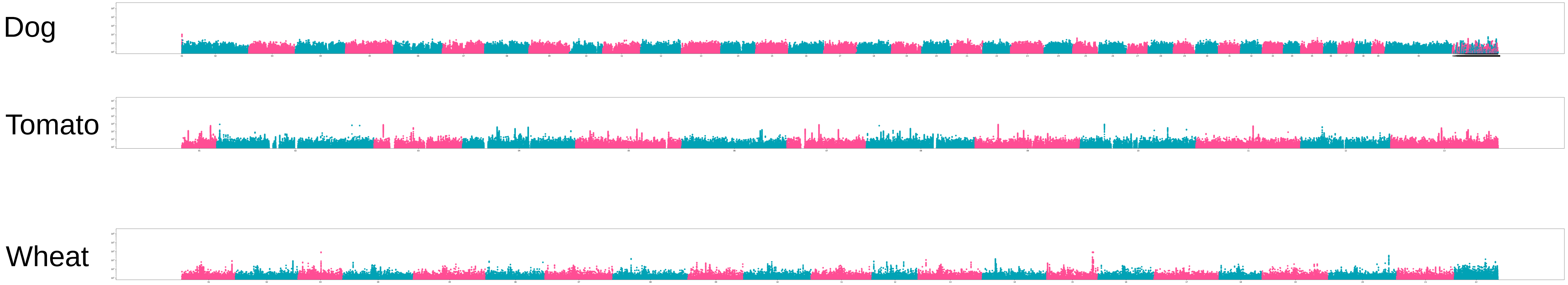

SFigure 3. Distribution of mapped fragments along genome

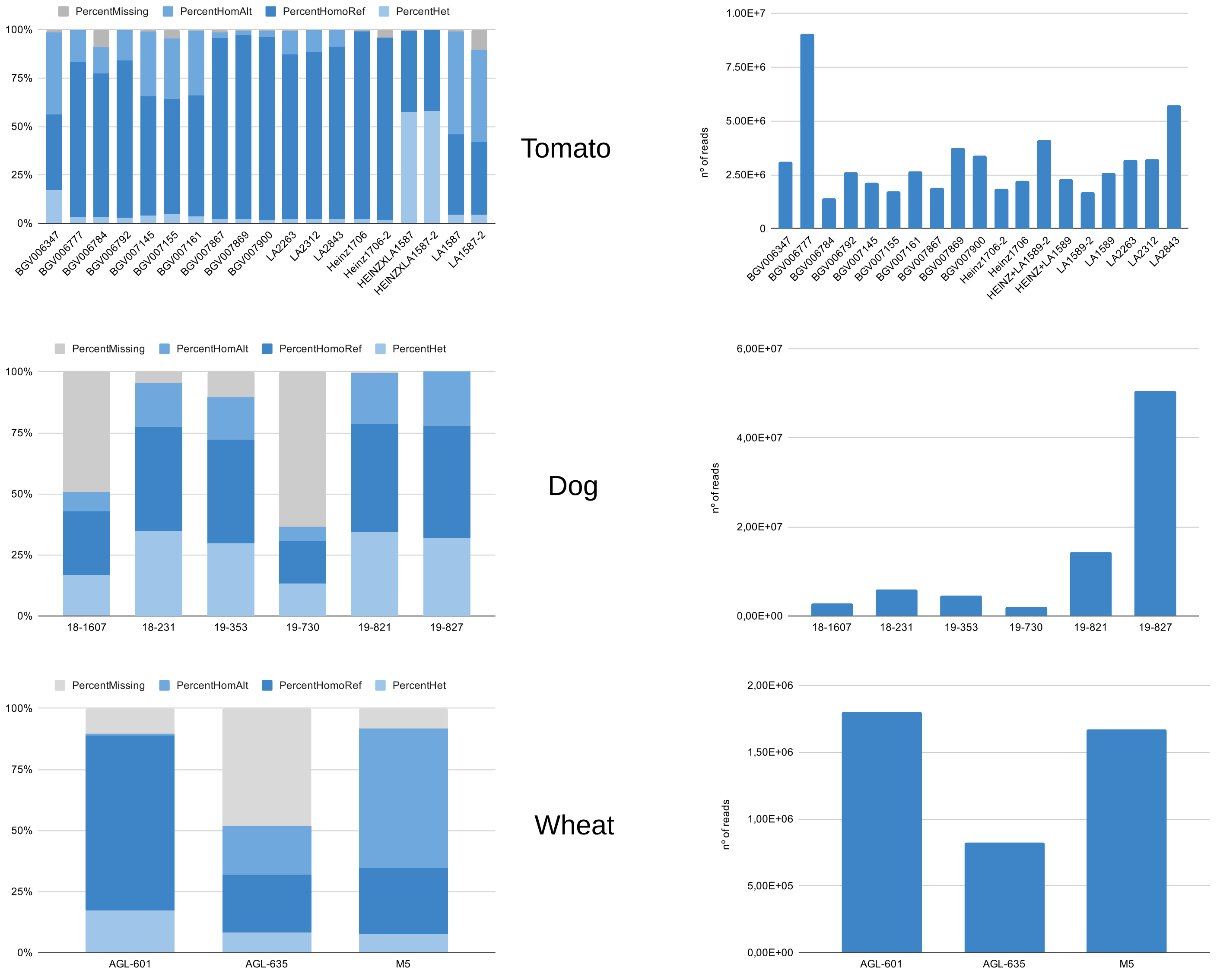

SFigure 4. SNP and mapping statistics

### Sup. Table 1 K-seq samples data

| Sample | Variety/code | Species | Genome | Ploidy | Sex | Genome sequence |
|---|---|---|---|---|---|---|
| AGL-601 | CHINESE SPRING | Triticum aestivum subsp vulgare | AABBDD | 6N | | 161010_Chinese_Spring_v1.0_pseudomolecules |
| AGL-635 | THATCHER | Triticum aestivum subsp vulgare | AABBDD | 6N | | 161010_Chinese_Spring_v1.0_pseudomolecules |
| M5 | BGE18908 | Triticum aestivum subsp speltoic | AABBDD | 6N | | 161010_Chinese_Spring_v1.0_pseudomolecules |
| AGL-001 | BGE047503 | Triticum turgidum subsp dicocco | AABB | 4N | | 161010_Chinese_Spring_v1.0_pseudomolecules |
| AGL-022 | BGE047513 | Triticum turgidum subsp turgidur | AABB | 4N | | 161010_Chinese_Spring_v1.0_pseudomolecules |
| SVEVO | SVEVO | Triticum turgidum subsp durum | AABB | 4N | | 161010_Chinese_Spring_v1.0_pseudomolecules |
| JG-1 | Seed give F. Vazque: | Triticum urartu | AA | 2N | | 161010_Chinese_Spring_v1.0_pseudomolecules |
| JG-9 | Seed give F. Vazque: | Triticum monococum subsp boe | AA | 2N | | 161010_Chinese_Spring_v1.0_pseudomolecules |
| 5 | Seed give F. Vazque: | Aegilops speltoides | BB | 2N | | 161010_Chinese_Spring_v1.0_pseudomolecules |
| JG-6 | Seed give F. Vazque: | Aegilops tauschii | DD | 2N | | 161010_Chinese_Spring_v1.0_pseudomolecules |
| Dulcinea | Dulcinea | Capsicum Annum | | 2N | | CA_000512255.2_ASM51225v2_genomic |
| Listada de gandia | Listada de gandia | Solanum melogena | | 2N | | Eggplant_V3_Chromosomes |
| Rudolph | Rudolph | Solanum tubersoum | | 4N | | GCF_000226075.1_SolTub_3.0_genomic |
| Petuniaxhibrida | | Petuniaxhibrida | | 2N | | Petunia_axillaris_v1.6.2_genome_HiC |
| Heinz1706 | Heinz1706 | Solanum lycopersicum | | 2N | | S_lycopersicum_chromosomes.2.50 |
| LA1589 | LA1589 | Solanum pimpinelifollium | | 2N | | S_lycopersicum_chromosomes.2.50 |
| BGV006777 | BGV006777 | Solanum lycopersicum var. cerasiforme | | 2N | | S_lycopersicum_chromosomes.2.50 |
| BGV006784 | BGV006784 | Solanum lycopersicum var. cerasiforme | | 2N | | S_lycopersicum_chromosomes.2.50 |
| BGV006792 | BGV006792 | Solanum lycopersicum var. cerasiforme | | 2N | | S_lycopersicum_chromosomes.2.50 |
| LA2263 | LA2263 | Solanum lycopersicum var. cerasiforme | | 2N | | S_lycopersicum_chromosomes.2.50 |
| LA2312 | LA2312 | Solanum lycopersicum var. cerasiforme | | 2N | | S_lycopersicum_chromosomes.2.50 |
| LA2843 | LA2843 | Solanum lycopersicum var. cerasiforme | | 2N | | S_lycopersicum_chromosomes.2.50 |
| BGV007867 | BGV007867 | Solanum lycopersicum var. cerasiforme | | 2N | | S_lycopersicum_chromosomes.2.50 |
| BGV007869 | BGV007869 | Solanum lycopersicum var. cerasiforme | | 2N | | S_lycopersicum_chromosomes.2.50 |
| BGV007900 | BGV007900 | Solanum lycopersicum var. cerasiforme | | 2N | | S_lycopersicum_chromosomes.2.50 |
| BGV007145 | BGV007145 | Solanum pimpinelifollium | | 2N | | S_lycopersicum_chromosomes.2.50 |
| BGV007155 | BGV007155 | Solanum pimpinelifollium | | 2N | | S_lycopersicum_chromosomes.2.50 |
| BGV007161 | BGV007161 | Solanum pimpinelifollium | | 2N | | S_lycopersicum_chromosomes.2.50 |
| BGV006347 | BGV006347 | Solanum pimpinelifollium | | 2N | | S_lycopersicum_chromosomes.2.50 |
| 18/231 | Poodle | Cannis lupus familares | | 2N | Female | GCF_000002285.3_CanFam3.1_genomic |
| 19/827 | Spaniel breton | Cannis lupus familares | | 2N | Male | GCF_000002285.3_CanFam3.1_genomic |
| 19/353 | Yorkshire terrier | Cannis lupus familares | | 2N | Female | GCF_000002285.3_CanFam3.1_genomic |
| 19/730 | mix | Cannis lupus familares | | 2N | Female | GCF_000002285.3_CanFam3.1_genomic |
| 19/821 | mix | Cannis lupus familares | | 2N | Male | GCF_000002285.3_CanFam3.1_genomic |
| 18/1607 | mix | Cannis lupus familares | | 2N | Male | GCF_000002285.3_CanFam3.1_genomic |

| Name_ID_Study | Name_ID_SRA | Breed/CommonNa | BioProject | BioSample | SRA | Sex |
|---|---|---|---|---|---|---|
| Alaskan Malamute | AlaskanMalamute( | Alaskan Malamute | PRJNA448733 | SAMN08872810 | SRR7107992 | F |
| American Cocker S | CockerSpanielAme | Cocker Spaniel (An | SRP108905 | SRR5664964 | SRR5664964 | M |
| Beagle 1 | CFA.117995 | Beagle | PRJNA263947 | SAMN06159681 | SRR7107976 | M |
| Bearded Collie 2 | BeardedCollie02 | Bearded Collie | PRJEB16012 | SAMEA4505492 | SRR7107609 | M |
| Belgian Malinois 4 | MA0163 | Belgian Malinois | PRJEB16012 | SAMEA104032048 | SRR7107526 | F |
| Black and Tan Coo | Coonhound01 | Black and Tan Coo | PRJNA263947 | SAMN04196853 | SRR7107925 | F |
| Boston Terrier | BostonTerrier01 | Boston Terrier | PRJNA448733 | SAMN08872906 | SRR7120145 | F |
| Boxer | Boxer01 | Boxer | PRJNA255370 | SAMN02921305 | SRR7107773 | F |
| Bull Terrier 1 | BT1021 | Bull Terrier | PRJEB16012 | SAMEA104125116 | SRR7107556 | F |
| Bull Terrier Miniat | BT007 | Miniature Bull Terr | PRJEB16012 | SAMEA4506897 | SRR7107631 | M |
| Bulldog | Bulldog01 | Bulldog | PRJNA288568 | SAMN03801654 | SRR2095477 | F |
| Chihuahua 1 | Chihuahua01 | Chihuahua | PRJNA288568 | SAMN03801656 | SRR2095478 | F |
| Chinese Crested 1 | ChineseCrested01 | Chinese Crested | PRJNA261736 | SAMN03075611 | SRR7107801 | M |
| Chow Chow 2 | ChowChow02 | Chow Chow | PRJNA288568 | SAMN03801658 | SRR2094392 | F |
| Coyote 1 | Coyote01 | Coyote | PRJNA255370 | SAMN02921301 | SRR7107770 | |
| Dachshund 1 | CFA.107835 | Dachshund | PRJNA263947 | SAMN06159670 | SRR7107965 | M |
| Dalmatian 2 | Dalmatian01 | Dalmatian | PRJNA448733 | SAMN08872976 | SRR7120157 | F |
| Doberman Pinsche | DO242 | Doberman Pinsche | PRJEB16012 | SAMEA4505489 | SRR7107606 | F |
| English Cocker Spa | CP003 | English Cocker Spa | PRJEB16012 | SAMEA4506900 | SRR7107632 | M |
| English Pointer 1 | EnglishPointer01 | Pointer (English) | PRJNA288568 | SAMN03801665 | SRR2094398 | M |
| English Springer Sp | EnglishSpringerSpa | English Springer Sp | PRJNA263947 | SAMN03580391 | SRR7107883 | M |
| German Shepherd | DS043 | German Shepherd | PRJEB16012 | SAMEA4506895 | SRR7107629 | F |
| German Wirehaire | GermanWirehaire( | German Wirehaire | PRJEB13468 | SAMEA3928144 | SRR7107585 | F |
| Golden Retriever 1 | 140447_S11 | Golden Retriever | PRJNA448733 | SAMN08873033 | SRR7120160 | M |
| Great Dane 1 | DD116 | Great Dane | PRJEB16012 | SAMEA104091557 | SRR7107534 | M |
| Grey Wolf (Canis L | WolfTibetan01 | Grey Wolf (Canis L | PRJNA448733 | SAMN03652997 | SRR7107906 | M |
| Grey Wolf 10 | Wolf19 | Grey Wolf | PRJNA255370 | SAMN02921311 | SRR7107777 | M |
| Greyhound 2 | Greyhound02 | Greyhound | PRJNA247491 | SAMN03067872 | SRR7107789 | M |
| Grossspitz | GS104 | Grossspitz | PRJEB16012 | SAMEA104105252 | SRR7107551 | F |
| Iberian Wolf (Canis | Wolf39 | Iberian Wolf (Canis | PRJNA318403 | SAMN04851099 | SRR7107942 | F |
| Irish Setter | IrishSetter01 | Irish Setter | PRJNA448733 | SAMN08873112 | SRR7120167 | F |
| Irish Water Spanie | IrishWaterSpaniel( | Irish Water Spanie | PRJNA448733 | SAMN08873115 | SRR7120168 | F |
| Italian Greyhound | ItalianGreyhound0 | Italian Greyhound | PRJNA288568 | SAMN03801673 | SRR2094401 | M |
| Jack Russell Terrie | JackRussellTerrier( | Jack Russell Terrie | PRJNA263947 | SAMN03580384 | SRR7107878 | M |
| Jindo | Jindo01 | Jindo | PRJDB2266 | SAMD00009664 | SRR7107521 | M |
| Labrador Retriever | 91317_S7 | Labrador Retriever | PRJNA448733 | SAMN08873148 | SRR7120181 | M |
| Miniature Poodle 2 | MiniaturePoodle01 | Miniature Poodle | PRJNA448733 | SAMN08873175 | SRR7120187 | F |
| MIX: Dachsund | CFA.118000 | MIX: Dachsund | PRJNA263947 | SAMN06159686 | SRR7107981 | M |
| MIX: Golden Retrie | GR1078 | MIX: Golden Retrie | PRJEB16012 | SAMEA104091572 | SRR7107549 | F |
| MIX: Kerry Blue Te | MIX_KerryBlueTer | MIX: Kerry Blue Te | PRJNA263947 | SAMN04196846 | SRR7107918 | M |
| MIX: Labrador Ret | LA2382 | MIX: Labrador Ret | PRJEB16012 | SAMEA104125075 | SRR7107552 | F |
| MIX: Mixed Breed | MI016 | MIX: Mixed Breed | PRJEB16012 | SAMEA4506893 | SRR7107627 | F |
| MIX: Siberian Husk | MixedBreed07 | MIX: Siberian Husk | PRJNA448733 | SAMN08873188 | SRR7120190 | F |
| Pekingese | Pekingese01 | Pekingese | PRJNA288568 | SAMN03801676 | SRR2095500 | F |
| Pomeranian | Pomeranian01 | Pomeranian | PRJEB16012 | SAMEA4506892 | SRR7107626 | M |
| Portuguese Poden | PortuguesePodeng | Portuguese Poden | PRJNA263947 | SAMN03580388 | SRR7107881 | M |
| Portuguese Water | PortugueseWaterD | Portuguese Water | PRJNA448733 | SAMN08873227 | SRR7120201 | M |
| Red Wolf 1 | Wolf25 | Red Wolf | PRJNA255370 | SAMN02921317 | SRR7107783 | F |
| Rottweiler 1 | 165414_S20 | Rottweiler | PRJNA448733 | SAMN08873243 | SRR7120204 | F |
| Saint Bernard 1 | SaintBernard01 | Saint Bernard | PRJNA288568 | SAMN03801685 | SRR2095502 | F |
| Saluki 1 | Saluki01 | Saluki | PRJNA288568 | SAMN03801686 | SRR2095503 | F |
| Samoyed 1 | Samoyed01 | Samoyed | PRJNA448733 | SAMN08873258 | SRR7120211 | F |
| Scottish Terrier 1 | ScottishTerrier01 | Scottish Terrier | PRJNA288568 | SAMN03801688 | SRR2094409 | F |
| Shiba Inu 1 | CFA.107839 | Shiba Inu | PRJNA263947 | SAMN05770194 | SRR7107955 | F |
| Siberian Husky 1 | SiberianHusky01 | Siberian Husky | PRJNA288568 | SAMN03801690 | SRR2095539 | F |
| Sloughi 1 | Sloughi02 | Sloughi | PRJEB16012 | SAMEA4506885 | SRR7107619 | M |

| | | | | | | |
|---|---|---|---|---|---|---|
| Spinone Italiano | CFA.109669 | Spinone Italiano | PRJNA263947 | SAMN06159677 | SRR7107972 | F |
| Standard Poodle 1 | CFA.107842 | Standard Poodle | PRJNA263947 | SAMN06159675 | SRR7107970 | F |
| Standard Schnauze | CFA.118001 | Standard Schnauze | PRJNA263947 | SAMN06159687 | SRR7107982 | M |
| Tibetan Terrier 2 | TibetanTerrier02 | Tibetan Terrier | PRJNA263947 | SAMN03580406 | SRR7107898 | M |
| Toy Poodle | ToyPoodle01 | Toy Poodle | PRJNA288568 | SAMN03801692 | SRR2095540 | F |
| West Highland Wh | WW558 | West Highland Wh | PRJEB16012 | SAMEA104091561 | SRR7107538 | F |
| Yorkshire Terrier 2 | PER00075 | Yorkshire Terrier | PRJNA448733 | SAMN08873470 | SRR7120258 | F |
| Yorkshire Terrier 3 | BAN00032 | Yorkshire Terrier | PRJNA448733 | SAMN08873449 | SRR7120237 | F |
| Yorkshire Terrier 3 | PER00204 | Yorkshire Terrier | PRJNA448733 | SAMN08873476 | SRR7120264 | F |
| Yorkshire Terrier 4 | BAN00041 | Yorkshire Terrier | PRJNA448733 | SAMN08873450 | SRR7120238 | M |
| Yorkshire Terrier 4 | PER00409 | Yorkshire Terrier | PRJNA448733 | SAMN08873490 | SRR7120278 | F |
| Standard Poodle 2 | StandardPoodle01 | Standard Poodle | PRJNA288568 | SAMN03801691 | SRR2095325 | F |
| Brittany | BrittanySpaniel01 | Brittany | PRJNA288568 | SAMN03801653 | SRR2094390 | M |

| SRA Run | bioproject | biosample | Accession Number | Species | Country |
|---|---|---|---|---|---|
| SRR12171193 | PRJNA644494 | SAMN09229607 | BGV006347-GBS | Solanum pimpine | Peru: Morropon: La Matanza |
| SRR12171275 | PRJNA644494 | SAMN09229618 | BGV006777-GBS | Solanum lycoper | Ecuador: Tena: Puerto Misahualli |
| SRR12171253 | PRJNA644494 | SAMN15045111 | BGV006784-GBS | Solanum lycopersicu | Ecuador |
| SRR12171238 | PRJNA644494 | SAMN09229620 | BGV006792-GBS | Solanum lycoper | Ecuador: Tena: Puerto Napo |
| SRR12171242 | PRJNA644494 | SAMN15045119 | BGV007145-GBS | Solanum pimpinellifol | Unknown |
| SRR12171271 | PRJNA644494 | SAMN09229648 | BGV007161-GBS | Solanum pimpine | Ecuador: Pedernales: Pedernales |
| SRR12171235 | PRJNA644494 | SAMN09229662 | BGV007867-GBS | Solanum lycopersicu | Mexico: Uman: Uman |
| SRR12171232 | PRJNA644494 | SAMN15045109 | BGV007869-GBS | Solanum lycoper | Mexico |
| SRR12171234 | PRJNA644494 | SAMN09229672 | BGV007900-GBS | Solanum lycoper | Mexico: Huauchinango: Huauchinango |
| SRR12171265 | PRJNA644494 | SAMN15045106 | HEINZ1706-1-GE | Solanum lycoper | USA |
| SRR12171264 | PRJNA644494 | SAMN15045106 | HEINZ1706-2-GE | Solanum lycoper | USA |
| SRR12171212 | PRJNA644494 | SAMN15472363 | HEINZ1706+LA1 | Solanum | Unknown |
| SRR12171303 | PRJNA644494 | SAMN15472363 | HEINZ1706+LA1 | Solanum | Unknown |
| SRR12181073 | PRJNA644494 | SAMN15488734 | LA1589-1-GBS | Solanum pimpine | Peru |
| SRR12181072 | PRJNA644494 | SAMN15488734 | LA1589-1-GBS | Solanum pimpine | Peru |
| SRR12182112 | PRJNA644494 | SAMN15045108 | BGV007155-GBS | Solanum pimpine | Ecuador |
| SRR12182111 | PRJNA644494 | SAMN15045107 | LA2312-GBS | Solanum lycoper | Peru |
| SRR12182110 | PRJNA644494 | SAMN15045118 | LA2843-GBS | Solanum lycoper | Peru |
| SRR12182109 | PRJNA644494 | SAMN15045114 | LA2263-GBS | Solanum lycoper | Peru |

Sup. table 4. Statistics predicted regions

| | Dog | Tomato | Wheat |
|---|---|---|---|
| Genome size | 2,410,976,875 | 824,674,700 | 14,547,261,565 |
| Prediction size | 6,038,783 | 1,022,464 | 1,534,057 |
| Coverage 10X size | 19,368,633 | 3,708,530 | 920964 |
| % genome predicted | 0.25% | 0.12% | 0.01% |
| %genome coverage 10X | 0.80% | 0.45% | 0.006% |
| % gene genome | 50.51% | 13.19% | 2.33% |
| % gene prediction | 53.0% | 20.98% | 0.40% |
| % gene coverage 10X | 49.23% | 28.24% | 2.8% |
| % repetitive genome | 42.83% | 64.03% | 43.65% |
| %repetitive prediction | 76.4% | 72.2% | 23.10% |
| % repetitive coverage 10X | 55.33% | 62.71% | 22.56% |
| % prediction coverage 10X | 3.60% | 4.06% | 2.28% |

### Sup. Table 5. Reads and Maping statisitics

| Tomato | | | | | | LIBRARY STATS | | | | | | | | | | |
|---|---|---|---|---|---|---|---|---|---|---|---|---|---|---|---|---|
| | Reas_stats | | | | | bam stats no filter | | | | | | | Bam stats >57 | | | |
| library | raw_reads | %raw_reads_q>30 | | reads_in_bam | mapped_reads | %mapped | properly_paired | %properly | %duplicated_reads | | reads_in_bam | mapped_reads | %mapped | properly_paired | %properly | %duplicated_reads |
| BGV006347 | 3109146 | 96.9 | | 3109146 | 3053905 | 98.2 | 2848552 | 91.6 | 0 | | 2124332 | 2124332 | 68.3% | 2082355 | 67.0% | 0 |
| BGV006777 | 9049310 | 97.5 | | 9049310 | 8946723 | 98.9 | 8545940 | 94.4 | 0 | | 6917286 | 6917286 | 76.4% | 6807863 | 75.2% | 0 |
| BGV006784 | 1395390 | 96.5 | | 1395390 | 1368657 | 98.1 | 1282412 | 91.9 | 0 | | 1012081 | 1012081 | 72.5% | 989824 | 70.9% | 0 |
| BGV006792 | 2614570 | 97.3 | | 2614570 | 2583000 | 98.8 | 2436978 | 93.2 | 0 | | 1894988 | 1894988 | 72.5% | 1861290 | 71.2% | 0 |
| BGV007145 | 2136394 | 96.9 | | 2136394 | 2101147 | 98.4 | 1964098 | 91.9 | 0 | | 1619356 | 1619356 | 75.8% | 1588883 | 74.4% | 0 |
| BGV007155 | 1725058 | 96.8 | | 1725058 | 1693701 | 98.2 | 1595958 | 92.5 | 0 | | 1313285 | 1313285 | 76.1% | 1286524 | 74.6% | 0 |
| BGV007161 | 2684288 | 96.3 | | 2684288 | 2639899 | 98.3 | 2488608 | 92.7 | 0 | | 1963881 | 1963881 | 73.2% | 1926893 | 71.8% | 0 |
| BGV007867 | 1890570 | 96.3 | | 1890570 | 1859429 | 98.4 | 1750774 | 92.6 | 0 | | 1485239 | 1485239 | 78.6% | 1450157 | 76.7% | 0 |
| BGV007869 | 3762642 | 97.1 | | 3762642 | 3723856 | 99 | 3527674 | 93.8 | 0 | | 2855539 | 2855539 | 75.9% | 2796573 | 74.3% | 0 |
| BGV007900 | 3400540 | 96.6 | | 3400540 | 3360606 | 98.8 | 3181976 | 93.6 | 0 | | 2604695 | 2604695 | 76.6% | 2552156 | 75.1% | 0 |
| Heinz1706-2 | 1837322 | 96.6 | | 1837322 | 1800028 | 98 | 1701162 | 92.6 | 0 | | 1379463 | 1379463 | 75.1% | 1342483 | 73.1% | 0 |
| Heinz1706 | 2235912 | 97.1 | | 2235912 | 2212312 | 98.9 | 2099146 | 93.9 | 0 | | 1749362 | 1749362 | 78.2% | 1713238 | 76.6% | 0 |
| HEINZ+LA1589-2 | 4122630 | 97.5 | | 4122630 | 4063211 | 98.6 | 3820806 | 92.7 | 0 | | 2929913 | 2929913 | 71.1% | 2872473 | 69.7% | 0 |
| HEINZ+LA1589 | 2292108 | 96.9 | | 2292108 | 2258330 | 98.5 | 2112176 | 92.1 | 0 | | 1658706 | 1658706 | 72.4% | 1623330 | 70.8% | 0 |
| LA1589-2 | 1688210 | 96.6 | | 1688210 | 1650068 | 97.7 | 1539184 | 91.2 | 0 | | 1132858 | 1132858 | 67.1% | 1105965 | 65.5% | 0 |
| LA1589 | 2597392 | 96.1 | | 2597392 | 2548188 | 98.1 | 2364862 | 91 | 0 | | 1782020 | 1782020 | 68.6% | 1743175 | 67.1% | 0 |
| LA2263 | 3207116 | 96.5 | | 3207116 | 3163072 | 98.6 | 2932696 | 91.4 | 0 | | 2344777 | 2344777 | 73.1% | 2297594 | 71.6% | 0 |
| LA2312 | 3236352 | 97.2 | | 3236352 | 3201903 | 98.9 | 3049030 | 94.2 | 0 | | 2525399 | 2525399 | 78.0% | 2481945 | 76.7% | 0 |
| LA2843 | 5754286 | 97.3 | | 5754286 | 5694567 | 99 | 5404532 | 93.9 | 0 | | 4385126 | 4385126 | 76.2% | 4310240 | 74.9% | 0 |

| Wheat | | | | | | LIBRARY STATS | | | | | | | | | | |
|---|---|---|---|---|---|---|---|---|---|---|---|---|---|---|---|---|
| | Reas_stats | | | | | bam stats no filter | | | | | | | Bam stats >57 | | | |
| sample | raw_reads | %raw_reads_q>30 | | reads_in_bam | mapped_reads | %mapped | properly_paired | %properly | %duplicated_reads | | reads_in_bam | mapped_reads | %mapped | properly_paired | %properly | %duplicated_reads |
| 5 | 10797584 | 97.1 | | 10786324 | 10522136 | 97.6 | 8778326 | 81.4 | 0.0 | | 756239 | 756239 | 7.0 | 720872 | 44018 | 0.0 |
| AGL-001 | 3895132 | 97.1 | | 3891220 | 3835473 | 98.6 | 3638440 | 93.5 | 0.0 | | 1165378 | 1165378 | 44103 | 1148901 | 43980 | 0.0 |
| AGL-022 | 2181276 | 97.2 | | 2179264 | 2144803 | 98.4 | 2052446 | 94.2 | 0.0 | | 711571 | 711571 | 32.7 | 702280 | 32.2 | 0.0 |
| AGL-601 | 1804646 | 97.1 | | 1802874 | 1778729 | 98.7 | 1727924 | 95.8 | 0.0 | | 699750 | 699750 | 38.8 | 692810 | 38.4 | 0.0 |
| AGL-635 | 822692 | 96.6 | | 821740 | 798957 | 97.2 | 759284 | 92.4 | 0.0 | | 275216 | 275216 | 33.5 | 271974 | 33.1 | 0.0 |
| JG-1 | 11750190 | 96.9 | | 11740210 | 11562407 | 98.5 | 10848940 | 92.4 | 0.0 | | 2003827 | 2003827 | 43847 | 1963156 | 44028 | 0.0 |
| JG-6 | 9634370 | 97.0 | | 9624976 | 9485547 | 98.6 | 9011216 | 93.6 | 0.0 | | 1922227 | 1922227 | 20.0 | 1889794 | 44001 | 0.0 |
| JG-9 | 11351450 | 97.3 | | 11342150 | 11176413 | 98.5 | 9997664 | 88.1 | 0.0 | | 1082686 | 1082686 | 43960 | 1045798 | 43870 | 0.0 |
| M5 | 1674292 | 97.4 | | 1672136 | 1648781 | 98.6 | 1589920 | 95.1 | 0.0 | | 624742 | 624742 | 37.4 | 617895 | 37.0 | 0.0 |
| SVEVO | 8837918 | 97.2 | | 8830708 | 8730903 | 98.9 | 8469834 | 95.9 | 0.0 | | 3059690 | 3059690 | 34.6 | 3029728 | 34.3 | 0.0 |

| Dog | | | | | | LIBRARY STATS | | | | | | | | | | |
|---|---|---|---|---|---|---|---|---|---|---|---|---|---|---|---|---|
| | Reas_stats | | | | | bam stats no filter | | | | | | | Bam stats >57 | | | |
| 18-1607 | 2917520 | 96.7 | | 2915624 | 2882978 | 98.8 | 2791176 | 95.7 | 0.0 | | 2700422 | 2700422 | 92.6 | 2637468 | 90.4 | 0.0 |
| 18-231 | 6006594 | 96.2 | | 6002540 | 5948047 | 99.0 | 5765142 | 96.0 | 0.0 | | 5492141 | 5492141 | 91.4 | 5382732 | 89.6 | 0.0 |
| 19-353 | 4562442 | 96.5 | | 4559602 | 4521292 | 99.1 | 4381804 | 96.0 | 0.0 | | 4170048 | 4170048 | 91.4 | 4087295 | 89.6 | 0.0 |
| 19-730 | 2006520 | 96.2 | | 2005176 | 1979984 | 98.7 | 1919324 | 95.7 | 0.0 | | 1831019 | 1831019 | 91.3 | 1794421 | 89.4 | 0.0 |
| 19-821 | 14336652 | 96.2 | | 14327874 | 14227315 | 99.2 | 13812776 | 96.3 | 0.0 | | 13151261 | 13151261 | 91.7 | 12911401 | 90.1 | 0.0 |
| 19-827 | 50522266 | 96.2 | | 50491188 | 50162687 | 99.3 | 48413780 | 95.8 | 0.0 | | 46829228 | 46829228 | 92.7 | 45708809 | 90.5 | 0.0 |

| Potato | | | | | | LIBRARY STATS | | | | | | | | | | |
|---|---|---|---|---|---|---|---|---|---|---|---|---|---|---|---|---|
| | Reas_stats | | | | | bam stats no filter | | | | | | | Bam stats >57 | | | |
| Rudolph | 2771878 | 97.0 | | 2766440 | 2678469 | 96.6 | 2453710 | 88.5 | 0.0 | | 1430893 | 1430893 | 51.6 | 1393103 | 50.3 | 0.0 |

| Eggplant | | | | | | LIBRARY STATS | | | | | | | | | | |
|---|---|---|---|---|---|---|---|---|---|---|---|---|---|---|---|---|
| | Reas_stats | | | | | bam stats no filter | | | | | | | Bam stats >57 | | | |
| Listada | 2334444 | 96.9 | | 2330440 | 2246368 | 96.2 | 2063334 | 88.4 | 0.0 | | 282126 | 1282126 | 54.9 | 1262873 | 54.1 | 0.0 |

| Pepper | | | | | | LIBRARY STATS | | | | | | | |
|---|---|---|---|---|---|---|---|---|---|---|---|---|---|
| | Reas_stats | | | | | bam stats no filter | | | | | | | Bam stats >57 |

#### Sup. Table 5. Reads and Maping statisitics

| | | | | | | | | | | | | | | | | | |
|---|---|---|---|---|---|---|---|---|---|---|---|---|---|---|---|---|---|
| Dulcinea | 1355328 | 97.4 | | 1351854 | 1321858 | 97.5 | 1254046 | 92.5 | 0.0 | | | 1003401 | 1003401 | 74.0 | 989987 | 73.0 | 0.0 |
| Petunia | | | | | | LIBRARY STATS | | | | | | | | | | | |
| Reas_stats | | | | bam stats no filter | | | | | | | Bam stats >57 | | | | | | |
| Petunia-hibrida | 1683972 | 96.9 | | 1677882 | 1642380 | 97.9 | 1507054 | 89.8 | 0.0 | | 722871 | 722871 | 43.1 | 702592 | 41.9 | 0 | |

**Supplemental data 1. Scheme algorithm**

1. Identify all k-mers present on the genome

2. Annotate the positions of k-mers

3. Filter out k-mers

    3.1. with bad dust score
    3.2. with GC_content <35% or GC_content >57%

4. Select the 1000 best k-mers (By default we sort the kmers by abundance in non repetitive regions)

5. Filter kmer that does not keep the with primer3 filtering_criteria:

    5.1. MELTING_TEMPERATURE_THRESHOLD = 0
    5.2. MAX_COMP_END = 3
    5.3. MAX_COMP_ANY = 5
    5.4. MAX_COMP_ANY_FACTOR = 0.44
    5.5. "max_diff_temp": 40

6. Filter out reverse complementary k-mers

7. Select groups of 10 kmers compatibles between them using primer3

8. For each group of 10 kmer predict the products generated by all combination of pairs within a group

    8.1. Filter by correct strandness
    8.2. Filter by minimum and maximum length

9. Generate the statistics for each k-mer combination

#### Supplemental data 2. K-seq protocol

**1- First synthesis**

1 µl of each forward short-primers (5µM each)
2 µl dNTPs (25mM)
4,5 µl klenow buffer 10X
200-600 ng genomic DNA
$H_20$ until 45 µl final volume

<u>Reaction:</u>
4 min 95ºC
5 min 37ºC—add 1 µl Klenow (5U/µl Klenow Fragment (3'→5' exo)
20 min 75ºC

**2- First Exonuclease digestion**

+ 2µl Exonuclease I (E. coli) 20U/µl

<u>Reaction:</u>
60 min 37ºC
20 min 80ºC

**3- Second synthesis**

+1µl each reverse short-primers (5µM each)
+0.5 buffer Klenow 10X
+ H20 to 4 µl final volume

<u>Reaction:</u>
4 min 95ºC
5 min 37ºC. In this step add 1 µl Klenow (5U/µl)
20 min 75ºC

**4- Second Exonuclease digestion**

+ 1µl Exonuclease I (E. coli) 20U/µl

<u>Reaction:</u>
60 min 37ºC
20 min 80ºC

**5- PCR**

15 µl reaction 4
25µl Taq NEB 2X (M0270L Taq 2X Master MixNEB )
1 µl IDT-NXT adapter i7 (2.5µM)
1 µl IDT-NXT adapter i5 (2.5µM)
8µl $H_20$

<u>Reaction:</u>
1 min 95ºC
X cycles (usually between 10 and 15 cycles. Adjust cycle number to avoid high molecular weight band smear in agarose gel electrophoresis)
30s 95ºC
20s 62ºC
30s 68ºC
Last extension step. 5 min 68ºC

### Supplemental data 3. Primer sequences

**Primers K-seq**

Wheat

```
adaptF-9W    TCG TCG GCA GCG TCA GAT GTG TAT AAG AGA CAG NNN GAC GTA TCA
adaptR-3W    GTC TCG TGG GCT CGG AGA TGT GTA TAA GAG ACA GNN NTG TTC ATC C
adaptR-8W    GTC TCG TGG GCT CGG AGA TGT GTA TAA GAG ACA GNN NTT CCT CTT C
```

Dog

```
adaptF-2D    TCG TCG GCA GCG TCA GAT GTG TAT AAG AGA CAG NNN GAG AGG CAG
adaptR-6D    GTC TCG TGG GCT CGG AGA TGT GTA TAA GAG ACA GNN NAA GGG AAG A
adaptR-9D    GTC TCG TGG GCT CGG AGA TGT GTA TAA GAG ACA GNN NCT CCT TTC C
```

Tomato

```
adaptF-4T    TCG TCG GCA GCG TCA GAT GTG TAT AAG AGA CAG NNN TCA TCT TC
adaptF-5T    TCG TCG GCA GCG TCA GAT GTG TAT AAG AGA CAG NNN CAA AGA AG
adaptR-5T    GTC TCG TGG GCT CGG AGA TGT GTA TAA GAG ACA GNN NTG TTG ATG
```

**Primers PCR**

```
IDT-8nt-NXT_i7_1     CAAGCAGAAGACGGCATACGAGATACGATCAGGTCTCGTGGGCTC*G*G
IDT-8nt-NXT_i7_2     CAAGCAGAAGACGGCATACGAGATTCGAGAGTGTCTCGTGGGCTC*G*G
IDT-8nt-NXT_i7_3     CAAGCAGAAGACGGCATACGAGATCTAGCTCAGTCTCGTGGGCTC*G*G
IDT-8nt-NXT_i7_4     CAAGCAGAAGACGGCATACGAGATATCGTCTCGTCTCGTGGGCTC*G*G
IDT-8nt-NXT_i7_5     CAAGCAGAAGACGGCATACGAGATTCGACAAGGTCTCGTGGGCTC*G*G
IDT-8nt-NXT_i7_6     CAAGCAGAAGACGGCATACGAGATCCTTGGAAGTCTCGTGGGCTC*G*G
IDT-8nt-NXT_i7_7     CAAGCAGAAGACGGCATACGAGATATCATGCGGTCTCGTGGGCTC*G*G
IDT-8nt-NXT_i7_8     CAAGCAGAAGACGGCATACGAGATTGTTCCGTGTCTCGTGGGCTC*G*G
IDT-8nt-NXT_i7_9     CAAGCAGAAGACGGCATACGAGATATTAGCCGGTCTCGTGGGCTC*G*G
IDT-8nt-NXT_i7_10    CAAGCAGAAGACGGCATACGAGATCGATCGATGTCTCGTGGGCTC*G*G
IDT-8nt-NXT_i7_11    CAAGCAGAAGACGGCATACGAGATGATCTTGCGTCTCGTGGGCTC*G*G
IDT-8nt-NXT_i7_12    CAAGCAGAAGACGGCATACGAGATAGGATAGCGTCTCGTGGGCTC*G*G
IDT-8nt-NXT_i5_1     AATGATACGGCGACCACCGAGATCTACACATATGCGCTCGTCGGCAGCG*T*C
IDT-8nt-NXT_i5_2     AATGATACGGCGACCACCGAGATCTACACTGGTACAGTCGTCGGCAGCG*T*C
IDT-8nt-NXT_i5_3     AATGATACGGCGACCACCGAGATCTACACAACCGTTCTCGTCGGCAGCG*T*C
IDT-8nt-NXT_i5_4     AATGATACGGCGACCACCGAGATCTACACTAACCGGTTCGTCGGCAGCG*T*C
IDT-8nt-NXT_i5_5     AATGATACGGCGACCACCGAGATCTACACGAACATCGTCGTCGGCAGCG*T*C
IDT-8nt-NXT_i5_6     AATGATACGGCGACCACCGAGATCTACACCCTTGTAGTCGTCGGCAGCG*T*C
IDT-8nt-NXT_i5_7     AATGATACGGCGACCACCGAGATCTACACTCAGGCTTTCGTCGGCAGCG*T*C
IDT-8nt-NXT_i5_8     AATGATACGGCGACCACCGAGATCTACACGTTCTCGTTCGTCGGCAGCG*T*C
```

nucleotides labelled with * have a phosphotioate bond to prevent primer degradation by the DNA polymerase
